# Many rice genes are differentially spliced between roots and shoots but cytokinin has minimal effect on splicing

**DOI:** 10.1101/293217

**Authors:** Nowlan H. Freese, April R. Estrada, Ivory C. Blakley, Jinjie Duan, Ann E. Loraine

**Author notes:** Equal contribution (Nowlan H. Freese, April R. Estrada). Corresponding author. Nowlan Freese, April R. Estrada, Ivory C. Blakley, Jinjie Duan, Ann E. Loraine, Corresponding author.

## Abstract

Alternatively spliced genes produce multiple spliced isoforms, called transcript variants. In differential alternative splicing, transcript variant abundance differs across sample types. Differential alternative splicing is common in animal systems and influences cellular development in many processes, but its extent and significance is not as well known in plants. To investigate alternative splicing in plants, we examined RNA-Seq data from rice seedlings. The data included three biological replicates per sample type, approximately 30 million sequence alignments per replicate, and four sample types: roots and shoots treated with exogenous cytokinin delivered hydroponically or a mock treatment. Cytokinin treatment triggered expression changes in thousands of genes but had negligible effect on splicing patterns. However, many genes were alternatively spliced between mock-treated roots and shoots, indicating that our methods were sufficiently sensitive to detect differential splicing in a data set. Quantitative fragment analysis of reverse transcriptase-PCR products made from newly prepared rice samples confirmed nine of ten differential splicing events between rice roots and shoots. Differential alternative splicing typically changed the relative abundance of splice variants that co-occurred in a data set. Analysis of a similar (but less deeply sequenced) RNA-Seq data set from *Arabidopsis* showed the same pattern. In both the *Arabidopsis* and rice RNA-Seq data sets, most genes annotated as alternatively spliced had small minor variant frequencies. Of splicing choices with abundant support for minor forms, most alternative splicing events were located within the protein-coding sequence and maintained the annotated reading frame. A tool for visualizing protein annotations in the context of genomic sequence (ProtAnnot) together with a genome browser (Integrated Genome Browser) were used to visualize and assess effects of differential splicing on gene function. In general, differentially spliced regions coincided with conserved regions in the encoded proteins, indicating that differential alternative splicing is likely to affect protein function between root and shoot tissue in rice.

## INTRODUCTION

Differential splicing of pre-mRNA transcripts, called alternative splicing, enables one gene to produce multiple transcript variants encoding different functions. Alternative splicing is an almost universal phenomenon in higher eukaryotes, occurring to varying degrees in every animal and plant genome examined to date (Kalsotra and Cooper, 2011; Reddy *et al*., 2013). In animals, differential expression of splice variants has been recruited as a regulatory mechanism in multiple processes, such as sex determination in invertebrates and neuronal differentiation in mammals (Kalsotra and Cooper, 2011; Salz, 2011; Barbosa-Morais *et al*., 2012).

In plants, less is known about the functional significance and patterns of alternative splicing. However, several trends are apparent. Genes involved in circadian regulation are highly alternatively spliced, often producing multiple splice variants that fluctuate in concert with day/night cycling along with overall transcript abundance (Filichkin *et al*., 2015b). The serine and arginine-rich (SR) family of RNA-binding, splicing regulatory proteins is greatly expanded compared to mammals and includes many plant-specific forms (Kalyna and Barta, 2004; Barbosa-Morais *et al*., 2006; Plass *et al*., 2008; Filichkin *et al*., 2015a). SR transcripts themselves are also highly alternatively spliced in plants, with the relative abundance of these alternative transcripts varying according to environmental stresses and hormones (Palusa *et al*., 2007; Gulledge *et al*., 2012; Filichkin *et al*., 2015a; Keller *et al*., 2016; Mei *et al*., 2017).

A growing body of evidence indicates that cell and tissue specific regulation of alternative splicing occurs in plants, but its significance and extent is not well established (Vitulo *et al*., 2014; Li *et al*., 2016a; Sun *et al*., 2018). We previously found through analysis of RNA-Seq data from *Arabidopsis* pollen that the relative abundance of splice variants was similar between leaves and pollen, despite the differences between the two tissues (Loraine *et al*., 2013). However, this latter analysis was limited by having just one biological replicate for pollen and only two biological replicates for leaves. A more comprehensive analysis of multiple *Arabidopsis* data sets found a high incidence of isoform switching, in which the identity of the most prevalent variant differs between sample types (Vaneechoutte *et al*., 2017). However, this splicing diversity may have arisen in part from the heterogeneity of the data sets used, which were produced using rapidly changing (and improving) sequencing technologies at different times by different groups.

In this study, we used a well-replicated RNA-Seq data set from rice to re-examine prevalence of alternative splicing between tissues and hormone (cytokinin) treatment. This data set was previously generated to investigate cytokinin regulation of gene expression in roots and shoots from 10-day old rice seedlings (Raines *et al*., 2016). A parallel study produced an analogous data set from *Arabidopsis* for comparison, but was less deeply sequenced (Zubo *et al*., 2017). Both the rice and *Arabidopsis* RNA-Seq data sets included three biological replicates per sample type and four sample types – roots and shoots treated with exogenous cytokinin or a mock, vehicle-only treatment. In both data sets, the treatment triggered differential expression of thousands of genes, with roots affected to a greater degree than shoots.

For most alternatively spliced genes, regardless of whether or not they were differentially spliced, the relative abundance of splicing forms was highly skewed, with most alternatively spliced genes producing one major isoform. Nonetheless, there was a large minority of alternatively spliced genes where minor isoforms were more abundant and therefore seemed likely to affect gene function. We found that the relative abundance of splice variants for most alternatively spliced genes was remarkably stable, with very few differentially spliced genes between cytokinin treated and control samples. By contrast, many more genes were differentially spliced between roots and shoot, and most differential splicing occurred within the protein-coding sequence. Moreover, nearly every differential splicing event detected merely change the relative abundance of splice variants that co-occurred in the same sample. These results suggest differential alternative splicing likely contributes to gene function diversification between roots and shoots by moderating the relative abundance of splice co-expressed splice variants, but alternative splicing plays little role in cytokinin signaling.

## MATERIALS & METHODS

### RNA-Seq library preparation and sequencing

Rice and *Arabidopsis* samples were prepared and sequenced as described in (Raines *et al*., 2016; Zubo *et al*., 2017). Rice seedlings (Nipponbare) were grown hydroponically for ten days in a growth chamber set to 14 hours light (28°C) and 8 hours of dark (23°C) with light intensity 700 mmol m^-2^ s^-1^. Around six to ten seedlings were grown in the same pot, in four pots. On the tenth day, culture media was replaced with new media containing 5 μM of the cytokinin benzyladenine (BA) or 0.05 mN NaOH as a control. After 120 minutes, roots and shoots were harvested separately. Roots and shoots from treatment or control pots were pooled to form three replicates per treatment. RNA was extracted and used to synthesize twelve libraries from BA-treated and mock-treated roots and shoots. Libraries were sequenced on an Illumina HiSeq instrument for 100 cycles, yielding 100 base, single end reads. Sequence data are available from the Sequence Read Archive under accession SRP04905. Aligned, processed data are available from the Oct. 2011 rice genome assembly IgbQuickload directorie at http://igbquickload.org/quickload/0_sativa_japonica_0ct_2011/.

*Arabidopsis* plants were grown vertically on 1 × Murashige and Skoog agar with 1% sucrose for ten days under continuous illumination in a temperature-controlled growth chamber (Zubo *et al*., 2017). BA was applied to ten-day-old plants by immersing seedlings in MS solution containing 5 μM BA dissolved using dimethyl sulfoxide (DMSO), or DMSO without BA as a control, and gently shaken for two hours. Plants were harvested, and shoots and roots were collected separately. Libraries were sequenced on an Illumina HiSeq 2000 instrument for 100×2 cycles, yielding 100 base, paired end reads, to a depth of around 10 million sequenced fragments per library. Sequence data are available from the Sequence Read Archive under SRP049054 (rice data) and SRP059384 *(Arabidopsis* data). Aligned, processed data are available from the TAIR10 (June 2009) *Arabidopsis* genome assembly IgbQuickload directories at http://lorainelab-quickload.scidas.org/rnaseq/.

### Data processing

Rice sequences were aligned onto the *O. sativa japonica* genome assembly Os-Nipponbare-Reference-IRGSP-1.0 (Kawahara *et al*., 2013) and *Arabidopsis* sequences were aligned onto the TAIR10 June 2009 release of the *Arabidopsis* genome (Lamesch *et al*., 2012) using TopHat (Kim *et al*., 2013) and BowTie2 (Langmead and Salzberg, 2012) with maximum intron size set to 5,000 bases. A command-line, Java program called “FindJunctions” was used to identify exon-exon junctions from gapped read alignments in the RNA-Seq data. FindJunctions produces BED format files containing junction features, and the score field of the BED file lists the number of read alignments that supported the junction. Only reads that aligned to a unique location in the genome were considered. Source code and compiled versions of FindJunctions are available from https://bitbucket.org/lorainelab/findjunctions.

### Identification of alternative splicing events and differential splicing

To date, there have been two major releases of *O. sativa japonica* gene models: the MSU7 gene set (Kawahara *et al*., 2013) and the RAP-Db gene set (Sakai *et al*., 2013). The two gene model sets contain mostly the same data, but the MSU7 gene models appear to be the most heavily used and annotated with Gene Ontology terms. For simplicity, and to take advantage of available functional annotations, we used the MSU7 annotations here. For analysis of *Arabidopsis* data, we used TAIR10 (Lamesch *et al*., 2012) and Araport11 (Cheng *et al*., 2017) gene models.

Annotated alternative splicing events and the number of reads supporting each alternative were identified using the exon-intron overlap method introduced in (English *et al*., 2010) and further developed here for use with RNA-Seq data. Exons and introns from pairs of gene models from the same locus were compared to identify alternatively spliced regions. Regions where an intron in one model overlapped an exon in another model on the same strand were identified and used to define mutually exclusive splicing choices (Fig. 1). Gene models that included an alternatively spliced region were designated the “L” form (for “Long”) with respect to the splicing choice. Likewise, models that lacked an alternatively spliced region were designated “S” (for “Short”). Alternatively spliced regions were labeled according to the type of alternative splicing, as follows. Regions flanked by alternative donor sites were designated “DS” for alternative <u>d</u>onor <u>s</u>ite. Regions flanked by alternative acceptor sites were labeled “AS” for alternative <u>a</u>cceptor <u>s</u>ite. AS and DS events that coincided with exon skipping were labeled “AS/ES” and “DS/ES”. Alternatively spliced regions arising from introns that the spliceosome sometimes failed to excise were designated “RI” for retained intron.

**Fig. 1.**
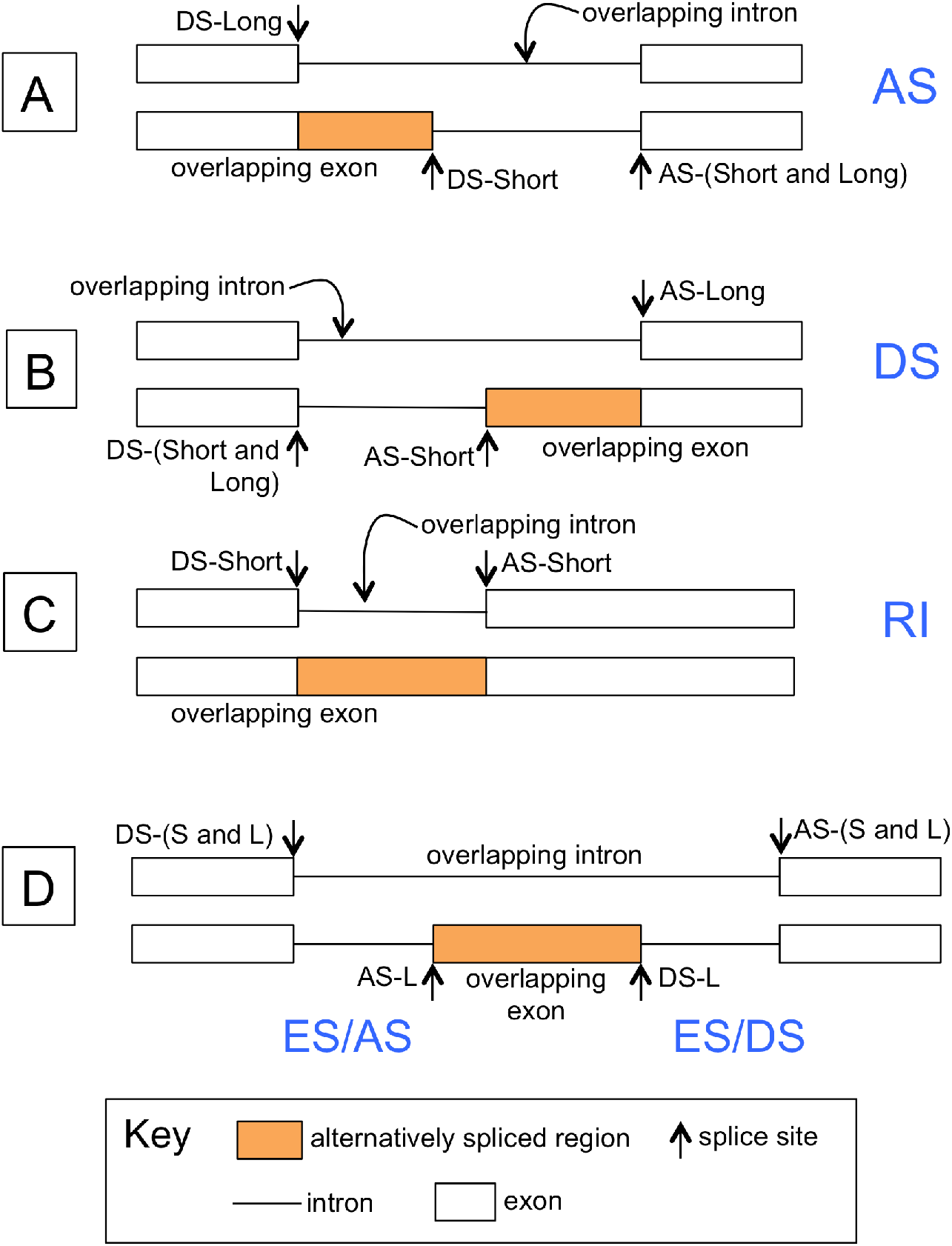
Alternative splicing annotation. The overlap between an intron in one gene model and an exon in another gene model defines an alternatively spliced region. Arrows indicate splice sites, named AS for acceptor site and DS for donor site. Use of sites named AS-L or DS-L causes inclusion of the differentially spliced region, generating the longer (L) isoform. Similarly, DS-S and AS-S refer to sites that exclude the differentially spliced region and generate the shorter (S) isoform. (**A**) Alternative donor sites, in which the U2 snRNP complex forms at alternative locations on the 5’ end of introns. (**B**) Alternative acceptor sites, in which the U1 snRNP complex forms at alternative sites near the 3’ end of alternatively spliced introns. (**C**) Alternatively spliced intron, in which a donor/acceptor site pairing can either be used or not. used, forming a retained intron (RI). (**D**) Alternatively spliced, skipped exon. In exon skipping, alternative splicing involves four sites, indicated by DS-S/L, AS-L, DS-L, and SD-S/L. Exon inclusion requires assembly of two spliceosome complexes linking DS-S/L with AS-L and DS-L with AS-S/L, while exon skipping requires linking DS-S/L and AS-S/L only.

For each alternatively spliced region representing two mutually exclusive splicing choices, RNA-Seq read alignments that unambiguously supported one or the other splicing choice were counted. For AS and DS events, only gapped reads that aligned across intron junctions were counted. For RI events, gapped reads that aligned across the retained intron were counted as support for the intron-removed (S) form, and un-gapped reads that overlapped at least 20 bases within the intron were counted as support for the intron-retained (L) form.

For each alternatively spliced region in each biological replicate, the number of reads supporting L or S, but not both, were used to calculate percent-spliced-in (PSI) as N/M*100, where N was the number of reads supporting the L form and M was the number of reads that supported S or L but not both. This is the same as the splicing index described in (Katz *et al*., 2010). To identify differentially spliced regions, a two-sided t-test was used to compare PSI between sample types. Because PSI variance was large for events with small M (very few informative reads), only alternatively spliced regions where M was 10 or more in at least three replicate libraries were tested. A false discovery rate (FDR) was calculated for each test using the method of Benjamini and Hochberg (Benjamini and Hochberg, 1995), as implemented in the R programming language “p.adjust” method. Alternative splicing events with FDR less than or equal to 0.1 were considered differentially alternatively spliced.

Software used to identify and quantify alternative events is available from https://bitbucket.org/lorainelab/altspliceanalysis. Data analysis code used to analyze RNA-Seq data is available from https://bitbucket.org/lorainelab/ricealtsplice. Data analysis code is implemented as R Markdown files designed to be run in the RStudio development environment. Readers interested in experimenting with different analysis parameters can clone the repository, modify the code, and re-run analyses as desired. RNA-Seq alignments, coverage graphs, and junctions data are available for visualization in Integrated Genome Browser (Freese *et al*., 2016).

### RT-PCR and capillary gel electrophoresis analysis of alternative splicing

Differential alternative splicing detected by analysis of RNA-Seq was re-tested using the reverse transcriptase, PCR-based fragment analysis method described in (Stamm *et al*., 2012). Differentially spliced regions identified computationally were PCR-amplified using fluorescently labeled primers and quantified using capillary gel electrophoresis. One benefit of the method is that the results are expressed as relative abundances of splice variants within a sample, thus eliminating the need to normalize using reference genes as in traditional qRT-PCR experiments aimed at measuring overall gene expression.

For splicing validation, new rice seedlings equivalent to the mock-treated (control) samples from the RNA-Seq experiment were grown and harvested. Seedlings were grown hydroponically in pots containing either liquid media only or calcined clay granules watered with liquid media as recommended in (Eddy *et al*., 2012). After twelve days, plants were removed from the pots and roots and shoots were collected separately. Roots and shoots from the same pot were combined to form paired biological replicates. Samples were flash frozen in nitrogen and stored at −80°C prior to RNA extraction.

RNA was extracted using the RNeasy Plant Mini Kit from Qiagen following the manufacturer’s instructions. First strand cDNA was synthesized using oligo dT primers and 1 μg of total RNA per 20 μL reaction. PCR amplification of cDNA was performed using primers flanking differentially spliced regions, including one primer labeled with 6-carboxyfluorescein (6-FAM) for amplicon detection during fragment analysis. Cycle parameters included denaturation at 94°C for 2 minutes, followed by 24 cycles of 94°C for 15 sec, 50°C for 30 sec and 70°C for 1 min, with a final elongation step of 72°C for 10 minutes. This was essentially the same regime described in (Stamm *et al*., 2012) but with fewer cycles to ensure reactions were stopped before exiting the logarithmic phase. PCR products were combined with size standards and separated on a 3730 Genetic Analyzer (Life Technologies). Amplicons were quantified using manufacturer-provided software by calculating the area under each amplicon peak. The percentage of the variant containing the alternatively spliced region (%L, see above) was calculated by dividing the long form area by the total area for both long and short forms. Spreadsheets with data exported from the instrument, along with PSI calculations, are available in the project git repository (https://bitbucket.org/lorainelab/ricealtsplice) in a subfolder named “Experimental Testing.”

## RESULTS

### Most genes annotated as alternatively spliced favored one dominant isoform

Using the exon-intron overlap method described previously (English *et al*., 2010) and Fig. 1, alternative splicing events within each gene were identified and annotated. Following annotation of alternative splicing events, RNA-Seq read alignments from the rice *and Arabidopsis* libraries described in (Raines *et al*., 2016; Zubo *et al*., 2017) were used to assess alternative splicing in four sample types: roots and shoots from seedlings treated with the cytokinin compound benzyladenine (BA) or with a mock, control treatment. For each alternative splicing event, the number of sequence alignments unambiguously supporting each alternative was counted. These counts were used to calculate percent-spliced-in (PSI), the percentage of read alignments supporting the longer (L) isoform.

In the combined data from all libraries from the rice data set, 77% of AS events had at least one read supporting each of the two splicing choices, and 19.8% had support for just one splicing choice. Only 2.8% of AS events has no reads supporting either form; these corresponded to genes with low or no expression in any of the sample types tested. Most genes annotated as alternatively spliced had small minor variant frequencies, i.e., the less frequently observed forms were supported by fewer than 20% of informative RNA-Seq sequences (Fig. 2). Nevertheless, there was a large minority of alternative splicing events (around one third) where the minor, less-frequently observed form was more abundant and was supported by at least two out of ten informative alignments. These alternatively spliced regions correspond to the middle, trough-like region of Fig. 2.

**Fig. 2.**
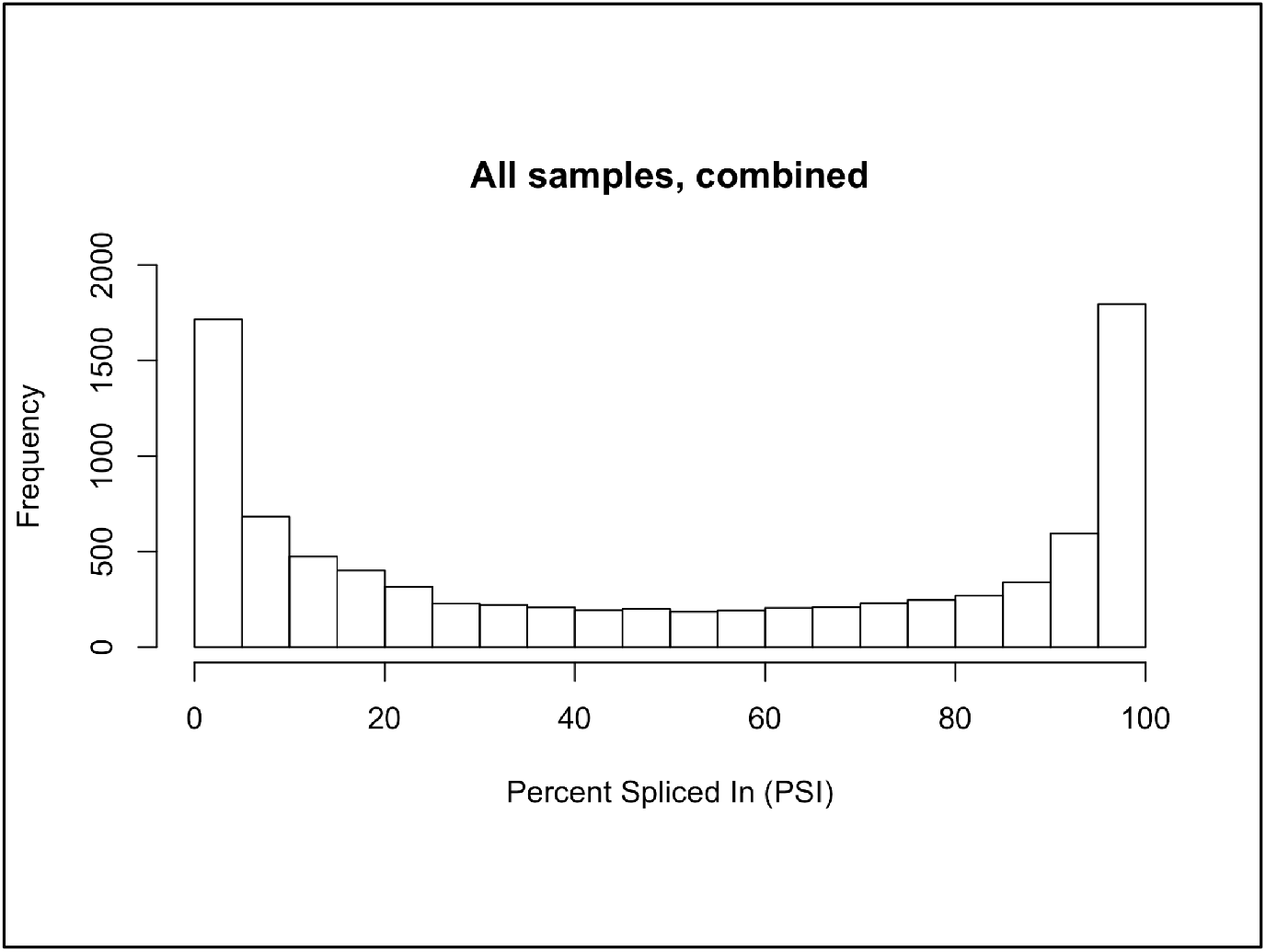
Distribution of percent-spliced-in (PSI) for annotated splicing events in rice where each choice was supported by at least one RNA-Seq alignment. PSI was calculated as 100*L/(S+L), where L and S were the number of reads that supported the splicing choice that included (L) or excluded (S) the differentially spliced region. Read alignment counts from all twelve libraries were combined to obtain a global view of alternative splicing occurrence in rice seedlings. The U-shaped character of the distribution persisted whether lower or higher thresholds of informative reads were used.

### Rice genes with abundant support for both alternative splicing choices perform many diverse functions

We used Gene Ontology term enrichment to determine if the subset of genes in rice for which alternatively spliced forms were unusually abundant exhibited enrichment with specific functions or processes, e.g., circadian cycling, in which alternative splicing might play a prominent regulatory role. We asked if some Gene Ontology terms were significantly enriched with genes containing alternative splicing events in which the minor form frequency was between 20 and 50%, corresponding to the central trough region of Fig. 2. Interestingly, we found that these genes exhibited a diversity of gene functions, with no significant enrichment of functional categories. Thus, alternative splicing in which minor forms are highly prevalent appears to affect genes with many functions in rice.

### Many rice genes are differentially spliced between roots and shoots but cytokinin hormone application has minimal effect on splicing

In animals, differential splicing between cell or tissue types contributes to cellular differentiation, especially in the nervous system (Naftelberg *et al*., 2015). Less is known about the role of alternative splicing in regulating cellular differentiation and other processes in plants. Rice shoots and roots are profoundly different tissues, but our previous analysis of this same data set found that many of the same genes were expressed in both (Raines *et al*., 2016). This raises the question of how these two different tissues are able to carry out their specialized roles, and suggest the hypothesis that differential splicing could enable differential functions in genes expressed in both tissues, as proposed in (Reddy *et al*., 2013).

Analysis of the effects of cytokinin treatment on this same data set from rice identified many thousands of genes that were differentially expressed in response to cytokinin (Raines *et al*., 2016). However, little is known about the role of alternative splicing during cytokinin response, except for one study in *Arabidopsis* that reported a shift in splicing of SR protein genes following cytokinin hormone treatment (Palusa *et al*., 2007). Therefore, we examined differential splicing in the rice RNA-Seq data set comparing root and shoot tissue with or without cytokinin.

First, we asked: When an alternatively spliced gene was expressed in two different sample types, was the relative abundance of splice variants the same or different? To address this, we examined correlation of PSI between roots and shoots or between BA-treated versus mock-treated samples (Fig. 3). We found that PSI was similar between treated and untreated samples, as revealed by the tighter clustering of scatter plot points (Fig. 3A-B). This indicated that genes that were alternatively spliced in BA-treated samples were also alternatively spliced in the controls, and that the relative abundance of splice variants was similar. Thus, the cytokinin hormone treatment had minimal effect on splicing. By contrast, there were many genes where the relative abundance of splice variants was different between roots and shoots (Fig. 3C).

**Fig. 3.**
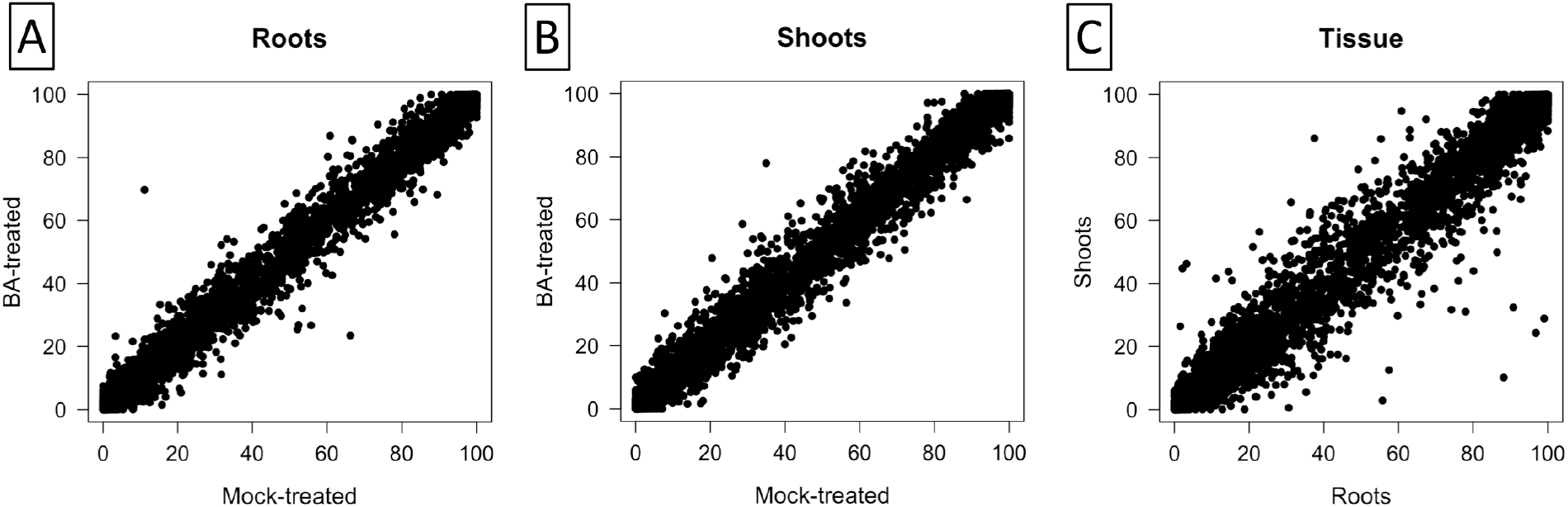
Scatter plots comparing percent-spliced-in (PSI) between sample types in rice for annotated splicing events. PSI was calculated from RNA-Seq reads obtained from sequencing rice seedling shoots and roots grown hydroponically and subjected to a two-hour treatment with BA, a cytokinin analog, or a mock-treatment (control). PSI is the average of three biological replicates. Only events with at least 15 informative read alignments in all six samples being compared were included. (A) BA-treated rice roots (y axis) compared to mock roots (x axis). (B) BA-treated rice shoots (y axis) compared to mock shoots (x axis). (C) Mock shoots (y axis) compared to mock roots (x axis).

Consistent with Fig. 3, statistical testing of PSI differences between sample types identified 90 genes where PSI was significantly different between roots and shoots (FDR ≤ 0.1) but only four and two genes where PSI was different between cytokinin-treated samples and controls in roots and shoots, respectively (Supplementary Table S1). Thus, we observed limited but non-trivial levels of differential alternative splicing between roots and shoots but minimal differential alternative splicing between control and BA-treated samples.

### Comparison of *Arabidopsis* differential splicing shows similar patterns to rice

To determine whether the observed patterns of differential splicing are similar in other plants, we analyzed splicing in *Arabidopsis* roots and shoots that had also been treated with cytokinin (Zubo *et al*., 2017). Due to the *Arabidopsis* libraries not being sequenced to the same depth as the rice libraries, many more splicing events had little or no support. Using the same FDR threshold as with the rice data set (FDR ≤ 0.1), we identified few differentially spliced regions between shoots and roots (3) and none in the control to treatment comparisons (Supplementary Table S2). However, PSI was distributed similarly to rice in that most alternatively spliced genes expressed one major isoform (Supplementary Fig. S1A). In addition, scatter plots showing average PSI in treated versus untreated samples showed a much tighter clustering of points as compared to scatter plots comparing roots and shoots (Supplementary Fig. S1B-D). Statistical testing of PSI differences confirmed the cytokinin hormone treatment had minimal effect on splicing in *Arabidopsis*. Thus, the general pattern of more differential splicing between tissue types as compared to treatment with exogenous cytokinin appears conserved between rice and *Arabidopsis*.

### Alternative splicing remodeled protein-coding sequence more often than disrupting it in rice

Alternative splicing can occur anywhere in a gene, including UTR and protein-coding regions. Most differential splicing between roots and shoots (67%) occurred within protein-coding regions (Table 1 and Supplementary Table S1), suggesting that differential splicing is likely to affect gene function at the level of the protein product. In every instance of differential alternative splicing, major and minor isoforms were both detected, with differential splicing observed as a change in the relative abundance of the two forms.

**Table 1.** *Counts of alternative splicing choices in rice that produce difference regions evenly divisible by three or with remainder of 1 or 2*.

| Alternative Splicing |  | Divisible | Remainder | Remainder | *P-value |
| --- | --- | --- | --- | --- | --- |
| Location | Event | by 3 | of 1 | of 2 |  |
| Coding region | Annotated as alternatively spliced | 3,248 | 2,466 | 2,411 | 3e-36 |
| UTR | Annotated as alternatively spliced | 1,152 | 1,127 | 1,113 | 1 |
| Coding region | Minor form is expressed | 173 | 149 | 153 | 8e-6 |
| UTR | Minor form is expressed | 34 | 20 | 13 | 0.03 |
| Coding region | Differentially spliced | 33 | 24 | 18 | 0.03 |
| UTR | Differentially spliced | 6 | 5 | 11 | 0.79 |

\*P-value obtained from binomial test of the null hypothesis that the true probability of a differentially spliced region having a length divisible by three is 1 in 3 and an alternative hypothesis that the probability is greater than 1 in 3.

Because three bases encode one amino acid, the lengths of spliced coding regions in a transcript are multiples of three. Thus, when alternatively spliced regions occur in coding regions and are not multiples of three, then inclusion of these regions in transcripts is likely to introduce a frame shift, resulting in a premature stop codon and a truncated protein product. As shown in Table 1, there was an enrichment of alternatively spliced regions in rice that were evenly divisible by three in coding regions versus non-coding in all subsets of the data. These subsets included all annotated alternatively spliced regions, regions where the minor form was unusually prevalent (the trough region of Fig. 2), and differentially spliced regions. Thus, alternative splicing within the coding regions of genes was biased against introducing frame shifts and promoted protein remodeling rather than truncation.

To further understand the effects of splicing on protein-coding sequences, we visualized differentially spliced regions together with RNA-Seq alignments, coverage graphs, and inferred junctions using genome browsers. Two genome browsers were used to visualize the data - Integrated Genome Browser (Freese *et al*., 2016) and ProtAnnot (Mall *et al*., 2016). Integrated Genome Browser (IGB) was used to examine RNA-Seq read alignments and compare alignments to the annotated gene structures. ProtAnnot, an IGB App, was used to search the InterPro database of conserved protein motifs to find out how (or if) splicing inferred from RNA-Seq data was likely to affect gene function through remodeling of protein motifs as detected by the InterProScan Web service (Finn *et al*., 2017).

Of the 105 differentially spliced regions, 71 overlapped protein-coding sequence regions, suggesting that in these cases, alternative splicing affected protein function. All but one (70/71) of the differentially spliced regions embedded in coding regions overlapped a predicted functional motif (e.g., a predicted transmembrane helix) or a region found by protein classification systems (e.g., Pfam (Finn *et al*., 2016) or PANTHER (Thomas *et al*., 2003)) to be conserved among members of the same protein family (Supplementary Table S1 and Fig. 4).

**Fig. 4.**
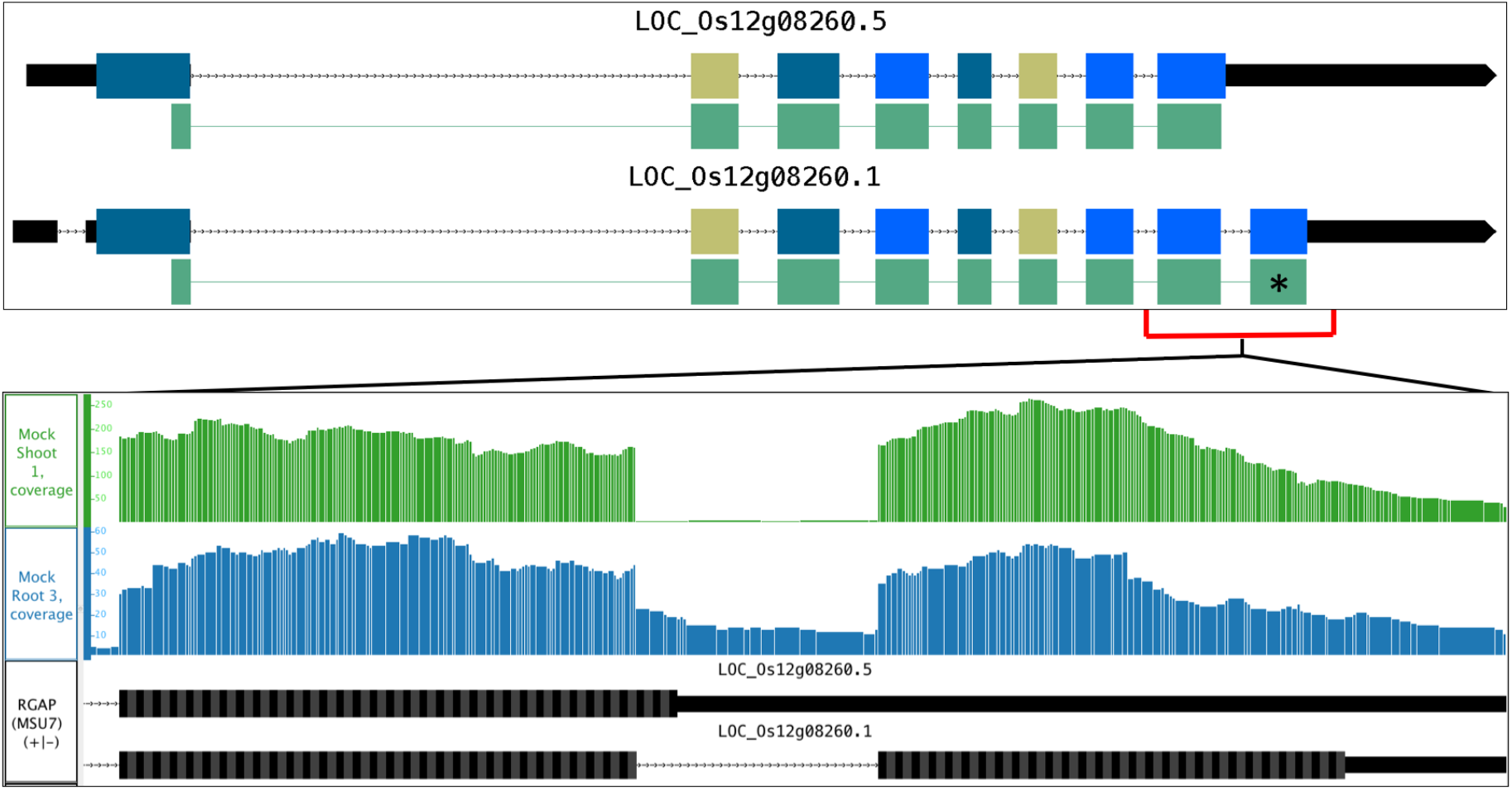
ProtAnnot and IGB images showing difference in splicing between rice shoot and root. ProtAnnot (upper panel) shows coding region exons color-coded by frame, with regions matching InterPro profiles indicated by green, linked rectangles. Asterisk highlights difference in the PANTHER InterPro profile PTHR11516 between isoforms 1 and 5 of the LOC_Os12g08260 gene. Integrated Genome Browser (lower panels) shows a zoomed-in view of RNA-Seq coverage graphs from rice root (blue) and shoot (green). Y-axis is the number of RNA-Seq aligned sequences with MSU7 gene models in black below.

### RT-PCR with capillary gel electrophoresis confirmed differential splicing between rice roots and shoots for nine of ten genes tested

We used a method based on capillary gel electrophoresis of fluorescently tagged PCR products to assay alternative splicing of ten genes detected as differentially spliced between rice roots and shoots (Stamm *et al*., 2012). New rice seedlings were grown under a close-to-identical replication of the RNA-Seq experiment. Primers were designed to amplify differentially spliced regions, including one primer that was conjugated to a fluorescent tag. Following PCR amplification of cDNA prepared from the new rice samples, products were resolved on a capillary-based sequencer and PSI calculated (Table 2). In nine out of ten genes, differential alternative splicing was confirmed. In the one case where differential alternative splicing was not confirmed, there were very few RNA-Seq read alignments covering the differentially spliced region, suggesting this was likely a false positive result. The FDR cutoff used to detect differential splicing in the RNA-Seq data was 0.1, corresponding to one in ten false discoveries, in line with results from the microcapillary-based analysis.

**Table 2.** *Differential splicing detected using RNA-Seq and re-tested using capillary gel electrophoresis (CGE) in rice*.

| Gene | AS type | Avg. RPKM Expression |  | RNA-Seq PSI (%L) |  | CGE PSI (%L) |  | *P-value |
| --- | --- | --- | --- | --- | --- | --- | --- | --- |
|  |  | Root | Shoot | Root | Shoot | Root | Shoot |  |
| LOC_Os01g25484, ferredoxin nitrite reductase | RI | 300 | 131 | 74.2 | 31.7 | 29.9 | 13 | 5e-05 |
| LOC_Os01g35580, unknown | AS | 55.7 | 34.9 | 44.0 | 66.3 | 49.9 | 66.9 | 2.47e-04 |
| LOC_Os01g45274, carbonic anhydrase | ES | 171 | 1,380 | 96.8 | 24.3 | 97.4 | 15.4 | 3e-09 |
| LOC_Os01g51290, protein kinase | RI | 49.6 | 49.9 | 88.4 | 95.1 | 13.3 | 17.2 | 0.03466 |
| LOC_Os03g05390, citrate transporter | RI | 219 | 174 | 86.0 | 96.3 | 86 | 95.5 | 1.2e-04 |
| LOC_Os12g08260, dehydrogenase E1 | RI | 8.03 | 30.3 | 55.7 | 2.9 | 4.1 | 0.87 | 3e-05 |
| LOC_Os01g61670, ureidoglycolate hydrolase | DS | 59 | 37.8 | 78.0 | 31.0 | 59 | 37.8 | 1.6e-09 |
| LOC_Os05g48040, MATE efflux family protein | DS | 7.05 | 6.72 | 88.1 | 100.0 | 89.4 | 88.27 | 0.632 |
| LOC_Os02g05830, ribulose biphosphate carboxylase | RI | 4.79 | 12.3 | 88.2 | 10.1 | 23.9 | 3.06 | 8e-04 |
| LOC_Os06g05110, superoxide dismutase | RI | 12.5 | 39.7 | 38.5 | 13.6 | 22.7 | 6.5 | 1.3e-07 |

\*P-value obtained from comparing roots and shoots PSI from CGE.

## DISCUSSION

This study profiled alternative splicing using a high coverage RNA-Seq data from 10-day old, hydroponically-grown rice seedlings treated with a cytokinin hormone. A less-deeply sequenced data set from similarly treated *Arabidopsis* seedlings provided comparison with another plant species. We found that cytokinin treatment induced very few splicing changes between treated and untreated controls. However, there were many differences in splicing between untreated roots and shoots, and most of these changed the protein coding region of genes.

Palusa et al. found that BA-treatment of *Arabidopsis* seedlings triggered splicing changes in multiple SR genes (Palusa *et al*., 2007), encoding RNA-binding proteins whose counterparts in metazoans regulated alternative splice site choice. Their study used PCR amplification of cDNA followed by agarose gel electrophoresis to detect changes in splicing and focused on SR protein genes only. Thus, we expected to observe changes in SR gene splicing due to the cytokinin treatment, leading to changes in splicing for many downstream genes. However, no such differential splicing was apparent in either RNA-Seq data set tested. It is possible that the differences in methodology used to measure splicing changes between the two studies (RNA-Seq versus visualization of PCR amplification of cDNA) could account for the differences in observations. However, close examination of SR splicing genes in our dataset revealed no significant differences.

One possible explanation for why the cytokinin treatment had minimal effect on splicing was that the treatment itself was ineffective. However, differential expression analysis showed that many genes were up- or down-regulated by the treatment in the two data sets tested – rice and *Arabidopsis* (Raines *et al*., 2016). Fewer genes were detected as differentially expressed in the *Arabidopsis* data set, most likely reflecting higher variability between biological replicates combined with lower sequencing depth as compared to the rice data set. Nevertheless, known cytokinin response genes were differentially regulated in both data sets, showing the cytokinin treatment penetrated plant cells and induced stereotypical cytokinin signaling without also triggering changes in splicing.

The relative lack of differential splicing between cytokinin-treated and mock-treated samples suggests that cytokinin signaling does not employ alternative splicing as a regulatory mechanism to the same degree as with other plant hormones, notably abscisic acid (ABA). ABA plays a role in perception and response to stresses, especially desiccation stress (Maia *et al*., 2014). ABA also plays a role in regulating splicing of SR45, an SR-like protein, and SR45 plays a role in regulating downstream splicing of multiple genes (Cruz *et al*., 2014). Thus, there is a clear connection between stress signaling and the plant stress hormone ABA.

By contrast, cytokinin signaling involves transfer of phosphate groups between successive elements of a phosphorelay signaling pathway culminating in phosophorylation-dependent activation of Myb-type transcription factor proteins called type B ARRs. Cytokinin treatment has no or little effect on transcription of type B ARRs, the key regulators of cytokinin signaling (Argyros *et al*., 2008; Kieber and Schaller, 2018). In addition, type B transcriptional regulators are not highly alternatively spliced. By contrast, a closely related family of similar genes encoding so-called “pseudo-response regulators” have similar sequence to type B ARRs and are highly alternatively spliced (James *et al*., 2012). These genes are involved in regulating the circadian clock and have nothing to do with cytokinin signaling.

Using the same methods and data set, we identified a relatively large number of genes in rice (90) that were differentially spliced genes between shoots and roots, and we validated nine of ten using fragment analysis of independently produced rice samples. This observation of differential splicing between roots and shoots is important for two key reasons. First, it shows that our data analysis methods can identify differential alternative splicing in a data set. In other words, the roots versus shoots comparison provided a positive control for differential splicing detection. Second, the detection of differential splicing between roots and shoots illuminates the function of alternative splicing in plant cells. For most of the differentially spliced regions, both forms were present, and the difference in relative abundance between forms was often slight, rarely more than five or ten percent (Supplementary Table S1). Our data supports the growing body of evidence that alternative splicing is cell, tissue, and stage specific in plants (Vitulo *et al*., 2014; Gupta *et al*., 2015; Li *et al*., 2016a; Sun *et al*., 2018), including in roots (Li *et al*., 2016b). It is likely that alternative splicing plays a role in fine-tuning gene function to meet the needs of different plant tissues or cell types where a gene is expressed.

We also examined the prevalence of alternative splicing, independent from differential splicing. That is, we used RNA-Seq read alignments to assess how often annotated alternative splice sites were used in our RNA-Set data sets. For most genes annotated as alternatively spliced, the minor form frequency was typically low, accounting for less than 20% of sequence read alignments across the differentially spliced region. Genes where minor form frequency exceeded 20% exhibited a diversity of functions. Thus, many diverse processes in rice involved alternatively spliced genes in which splice variants were expressed at levels likely to affect gene function in different ways.

A major limitation of this study was that we limited our analysis to annotated splice forms and did not attempt to form new transcript models based on the RNA-Seq data. This was done mainly because the libraries used were not strand-specific and attempts to assemble transcripts using transcript assembly tools led to incorrect fusion of neighboring genes and other artifacts (not shown). Future studies will therefore benefit greatly from using better library preparation protocols to simplify and streamline data analysis. Nonetheless, this analysis provides new insight into the role of alternative splicing in plant tissues and hormone response.

In conclusion, by analyzing the number of reads that supported different splice variants, we identified examples of differential splicing with confirmation by RT-PCR with capillary gel electrophoresis. There were 90 genes differentially spliced between rice root and shoot tissues, but only four between cytokinin-treated and non-treated samples. For most differential splicing events, the protein-coding regions were affected, strongly suggesting that differential splicing is playing a role in modulating gene function between roots and shoots.

---

**Supplementary Table S1.** Spreadsheet containing the rice mock root vs mock shoot, mock root vs treated root, and mock shoot vs treated shoot splicing data.

**Supplementary Table S2.** Spreadsheet containing the *Arabidopsis* mock root vs mock shoot splicing data.

## ACKNOWLEDGEMENTS

This material is based upon work supported by the National Science Foundation through the Postdoctoral Research Fellowship in Biology under Grant No.1523814 awarded to NHF and the Plant Genome Research Program award number IOS-1238051. NIH award 2R01GM103463-06A1 to AEL supports development of the IGB software.

---

**Fig. S1.**
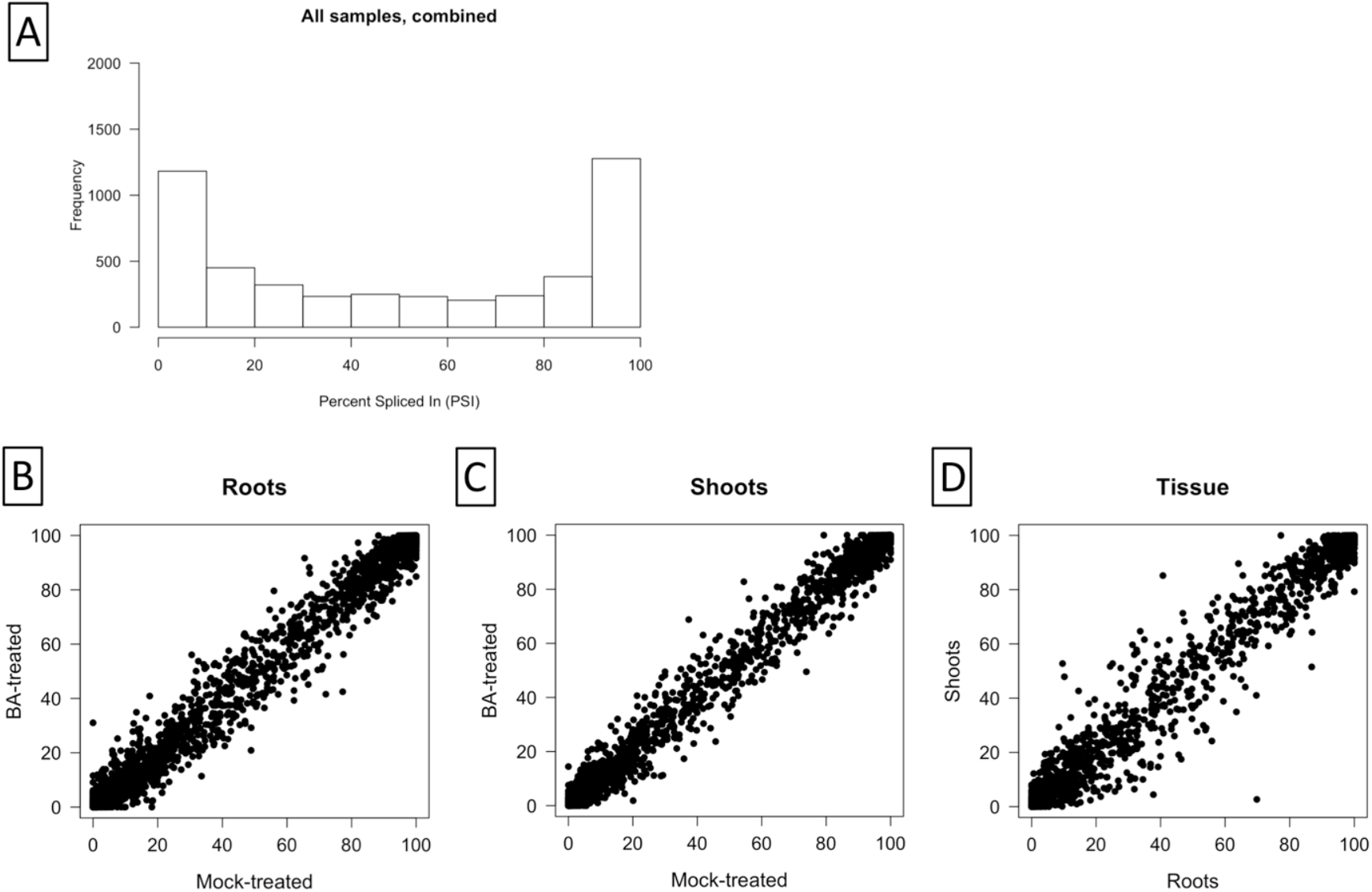
Percent-spliced-in (PSI) in *Arabidopsis* RNA-Seq data set between sample types for annotated splicing events. (A) Counts of PSI events in all samples combined. PSI was calculated from RNA-Seq reads obtained from sequencing *Arabidopsis* seedling shoots and roots grown with or without BA, a cytokinin analog. PSI is the average of three biological replicates. Only events with at least 10 informative read alignments in all six samples being compared were included. (B) BA-treated *Arabidopsis* roots (y axis) compared to mock roots (x axis). (C) BA-treated *Arabidopsis* shoots (y axis) compared to mock shoots (x axis). (D) Mock shoots (y axis) compared to mock roots (x axis).

## Abbreviations

SR: Serine/Arginine
BA: benzyladenine
L: long
S: short
FDR: false discovery rate
PSI: percent spliced in
AS: acceptor site
DS: donor site
ES: exon skipping
RI: retained intron
ABA: abscisic acid
IGB: Integrated Genome Browser
CGE: capillary gel electrophoresis
RPKM: Reads Per Kilobase per Million mapped reads

